# Virus-like particle capture reveals coordination of actin remodeling during Shigella flexneri entry by host proteins

**DOI:** 10.1101/2025.10.21.683715

**Authors:** María Luisa Gil-Marqués, Matthew S. Fish, Le Anh Thu Pham, Vritti Kharbanda, Keith T. Egger, Marcia B. Goldberg, Poyin Chen

**Affiliations:** Division of Infectious Diseases, Department of Medicine, Massachusetts General Hospital, Boston, Massachusetts, USA; Department of Microbiology, Blavatnik Institute, Harvard Medical School, Boston, Massachusetts, USA; Broad Institute, Cambridge, Massachusetts, USA

**Author notes:** Address correspondence to Poyin Chen. University of Massachusetts School of Medicine, Worcester, Massachusetts, USA.

**Keywords:** Shigella flexneri, type 3 secretion system, translocon, 14-3-3

## Abstract

*Shigella* spp. are intracellular bacterial pathogens that enter the host via plasma membrane insertion of a type 3 secretion system (T3SS) translocon, which triggers signaling cascades that include modulation of cytoskeletal dynamics, resulting in bacterial uptake. To better understand translocon insertion-induced host processes, we adapted a method to capture in virus-like particles (VLP) host proteins that are recruited to the cytosolic face of natively delivered *S. flexneri* translocons. Proteomic analyses reveal enrichment of 14-3-3ζ, a signaling protein, and CAP2, a regulator of actin turnover. 14-3-3ζ and CAP2 are necessary for host entry by T3SS pathogens. 14-3-3ζ dimers function as molecular scaffolds in the formation of bacterial-associated membrane ruffles. Concurrently, CAP2 localizes to membrane ruffles and cooperates with 14-3-3ζ to enable the formation of membrane ruffles that function efficiently in bacterial uptake. The findings define a coordinated role for 14-3-3ζ and CAP2 in cytoskeletal dynamics during T3SS pathogen infection.

## Introduction

*S. flexneri* is an intracellular bacterial pathogen that gains entry into host cells via the T3SS^1^. The T3SS, a conserved virulence strategy, encodes a nanomachine that delivers into the cytosol of host cells effector proteins that facilitate bacterial uptake into host cells and subsequent infection^2–6^. IpaB and IpaC, the first two proteins secreted through the T3SS upon *S. flexneri* contact with host cells, are translocases that form a heterooligomeric translocon in the plasma membrane. The T3SS translocon is highly conserved among T3SS pathogens and constitutes an early interface between the pathogen and the host cell^4,7,8^. The *S. flexneri* translocon interacts with host proteins and, in addition to serving as the conduit for effectors, controls their secretion from the bacterium into the cell^9–11^.

Plasma membrane insertion of translocons initiates a cascade of host responses that culminate in bacterial uptake by plasma membrane ruffling (micropinocytosis)^12^. Upon membrane insertion, the translocase IpaC interacts with host intermediate filaments, inducing conformational changes in the translocon that allow docking of the T3SS needle onto the pore^8,13^. Concurrently, actin polymerization is required to open the pore channel for delivery of effector proteins into the host cytosol^10^. Several *S. flexneri* T3SS effectors remodel the host cytoskeleton at the start of infection. IpaA binds to and activates host vinculin, triggering F-actin polymerization^14,15^, IpgB1 stimulates Rac1/Cdc42 to promote actin polymerization at entry sites, initiating membrane ruffle formation^16^, and VirA interacts with calpain to modulate bacterial entry^17^.

To extend our understanding of host processes that are triggered by translocon insertion and which contribute to bacterial entry, we identified host proteins that localize to sites of *S. flexneri* engagement with host plasma membranes. To do so, we captured in VLPs host proteins that accumulated at the cytosolic face of membrane-embedded translocons^18,19^. Proteomic analysis of these VLPs demonstrated enrichment of 14-3-3ζ, a protein involved in regulating cytoskeletal dynamics, at the cytosolic face of translocons. We found that upon *S. flexneri* engagement with host plasma membranes, 14-3-3ζ acts as a molecular scaffold, contributing to the expansion of the membrane ruffles that enable bacterial entry. Furthermore, we found that CAP2, an actin capping protein that promotes polymerization of branched actin networks, is enriched to *S. flexneri* membrane ruffles, dependent on 14-3-3ζ, and like 14-3-3ζ, is required for efficient *S. flexneri* entry.

## Results

### Adaptation of virus-like particle generation for capturing membrane-embedded *Shigella flexneri* translocons

To investigate host protein engagement with membrane-embedded *S. flexneri* translocons, we modified Virotrap, a method that employs HIV-1 Gag to capture plasma membrane and associated molecules within virus-like particles (VLPs) (Fig. 1a)^18,19^. Gag is sufficient to induce the budding of membrane-bound vesicles. HIV-1 Gag-transfected HeLa monolayers were infected with *S. flexneri* expressing a derivative of the translocase IpaB that is FLAG-tagged at the C-terminal extracellular domain.

**Fig. 1.**
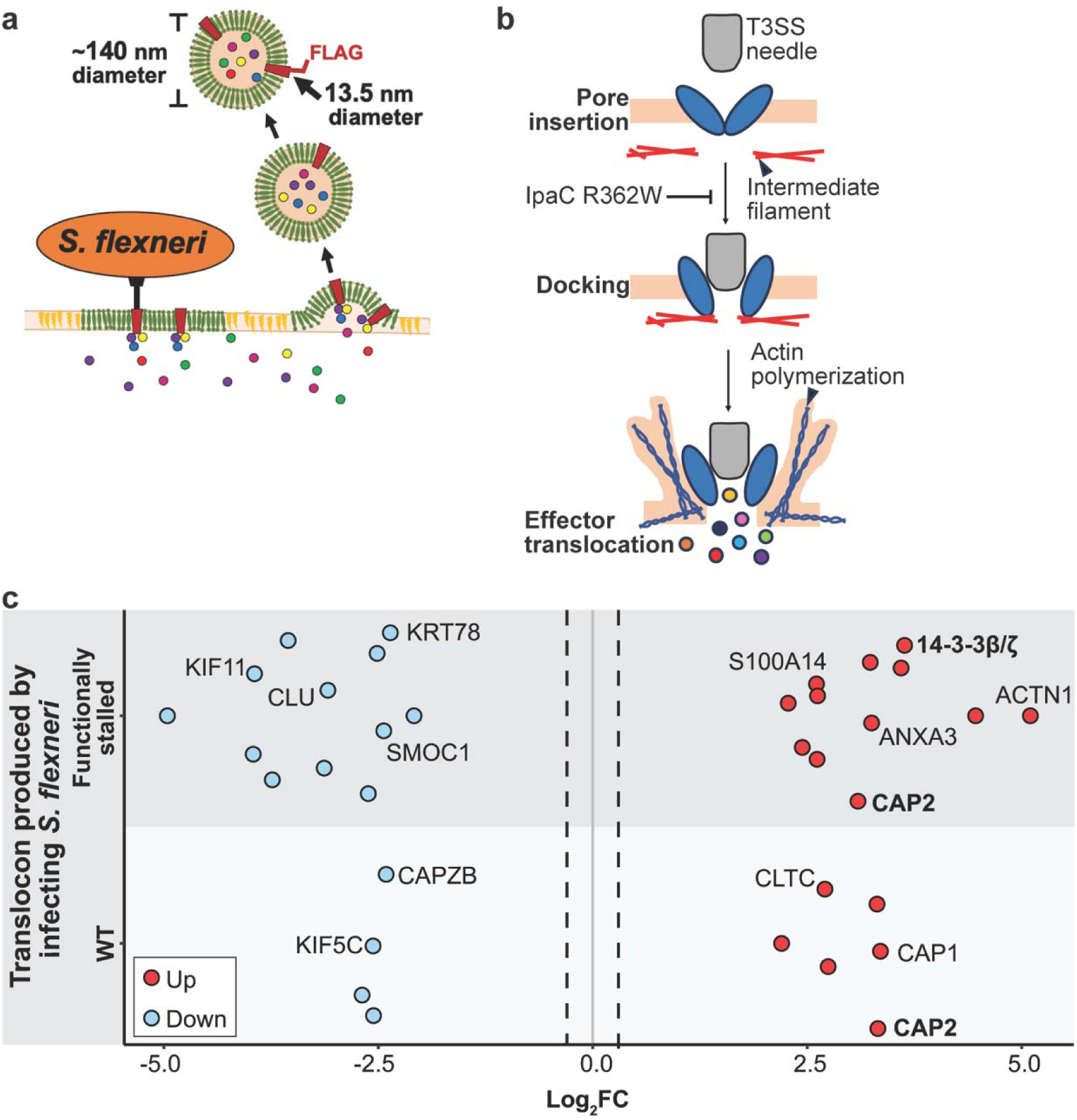
Proteomic analysis of VLPs that capture host factors proximal to type 3 translocons during *S. flexneri* invasion. **a,** Schematic of incorporation of *S. flexneri* type 3 translocons into VLPs. Following insertion of translocons into the plasma membrane, Gag-mediated membrane budding leads to incorporation of translocons into VLPs. *S. flexneri* translocase IpaB is tagged with FLAG at its C-terminus, which remains extracellular both following membrane insertion and on VLPs. The diameter of VLPs is approximately 140 nm, whereas that of translocons is approximately 13.5 nm. **b,** Schematic of type 3 translocons during cell invasion. The secretion system needle docks on plasma membrane-embedded translocons in a process that depends on interaction of IpaC with intermediate filaments^9^. Infection with *S. flexneri* expressing the IpaC derivative R362W, which is defective for interaction with intermediate filaments, leads to reduced bacterial docking but wildtype levels of translocon insertion^9^. Effector translocation depends on de novo actin polymerization^10^. **c,** Enrichment of cellular proteins in VLPs from HeLa cells infected with *S. flexneri* IpaC R362W (top) or WT *S. flexneri* (bottom). Mass spectrometry analysis of VLPs. Shown are significant (p ≤ 0.05) log_2_ fold-changes (Log_2_FC) in abundance of human proteins compared with uninfected cells. Labeled are cytoskeletal proteins, proteins involved in calcium signaling, and proteins known to be involved in *S. flexneri* infection. Bold, proteins investigated further in this study.

The presence of plasma membrane-localized Gag did not sterically hinder native delivery, membrane insertion, or function of translocon complexes (Extended Data Fig. 1). Assembly of translocons and effector delivery into Gag-transfected HeLa cells were comparable to those of untransfected cells (Extended Data Fig. 1a,b,c,d). VLPs containing FLAG-tagged translocons were selectively enriched from the culture supernatant of infected cells (Extended Data Fig. 1e). These data indicate that expression of Gag does not hinder *S. flexneri* translocon insertion, assembly, or function. Furthermore, this modified Virotrap approach enables targeted selection of translocon-containing VLPs for downstream analyses.

VLPs generated during *S. flexneri* infection were collected, and the subset that contained membrane-embedded translocons were enriched from the total VLP population via FLAG immunoprecipitation. VLP proteomes were analyzed by mass spectrometry.

### Cytoskeletal proteins are enriched at the cytosolic face of functionally stalled T3SS translocons

We collected in parallel VLPs containing functionally wildtype translocons and VLPs containing functionally stalled translocons. Stalled translocons were achieved by expressing *S. flexneri* IpaC R362W (Fig. 1b), a mutant IpaC derivative that locks the translocon in a docking incompetent state^13^. Stalled translocons theoretically promote accumulation of host proteins that participate in protein interactions at the cytosolic face of translocons^20^, as a result of delays in membrane ruffling and bacterial uptake. Collected VLPs were analyzed by mass spectrometry. IpaB and IpaC were detected in abundance (32-34 peptides and 10-12 peptides detected, respectively) in VLPs produced during *S. flexneri* infection compared to VLPs produced by uninfected HeLa monolayers, indicating the presence of translocons specifically in infection-associated VLPs. Also identified in infection-associated VLPs, but not in statistically significant abundances, were host cytoskeletal components known to be involved in *S. flexneri* infection, including actin (ACTA), vimentin, and spectrin (data available at the Proteomics Identifications Database, PRIDE), indicating that this approach is an effective method for capturing infection-associated host proteins.

When compared with VLPs produced from uninfected HeLa monolayers, VLPs containing functionally wildtype translocons displayed significantly increased abundance of cytoskeletal components (adenylyl cyclase-associated protein [CAP]1, CAP2), whereas VLPs containing functionally stalled translocons displayed significantly increased abundance of some of the same cytoskeletal components (CAP2), additional cytoskeletal components (ACTN1), and intracellular signaling proteins (ANXA3, S100A14, YWHAB [14-3-3β/ζ]) (Fig. 1c).

### 14-3-3ζ is required for efficient *S. flexneri* entry into host cells

14-3-3 proteins are present in all eukaryotic cell types and function as molecular scaffolds, facilitating numerous cellular processes and signaling pathways^21,22^. Their scaffolding functions depend on dimerization and binding to phosphoserine and phosphothreonine. In bacterial pathogenesis, 14-3-3 proteins are necessary for activation of *Pseudomonas aeruginosa* ExoS activity^23^.

Seven 14-3-3 isoforms exist, of which 14-3-3β and 14-3-3ζ share high sequence similarity^24,25^. In the proteome of VLPs from stalled translocons, the m/z ratios and corresponding sequences identified by mass spectrometry mapped to peptides present in both 14-3-3β and 14-3-3ζ, indicating that either or both isoforms might be associated with *S. flexneri* translocons. To discriminate which isoform was enriched by mass spectrometry, we generated HeLa cells lacking 14-3-3ζ. Loss of 14-3-3ζ significantly hindered bacterial adherence and entry. Compared to wildtype (WT) HeLa monolayers, *S. flexneri* entry into HeLa 14-3-3ζ^-/-^ monolayers was reduced by 50% ± 12 and 57% ± 5, as determined by gentamicin protection and plaque assays, respectively (Fig. 2a, Extended Data Fig. 3a,b), whereas bacterial cell-to-cell spread was not defective (Extended Data Fig. 3c). These data indicate that 14-3-3ζ is specifically required during *S. flexneri* entry and that 14-3-3β is non-redundant for this function. They also suggest that 14-3-3ζ is the isoform enriched in the VLPs from stalled translocons. Complementation of 14-3-3ζ rescued the bacterial entry and adherence phenotypes (Fig. 2a, Extended Data Fig. 3a). These results indicate that 14-3-3ζ is required for *S. flexneri* entry but not spread.

**Fig. 2.**
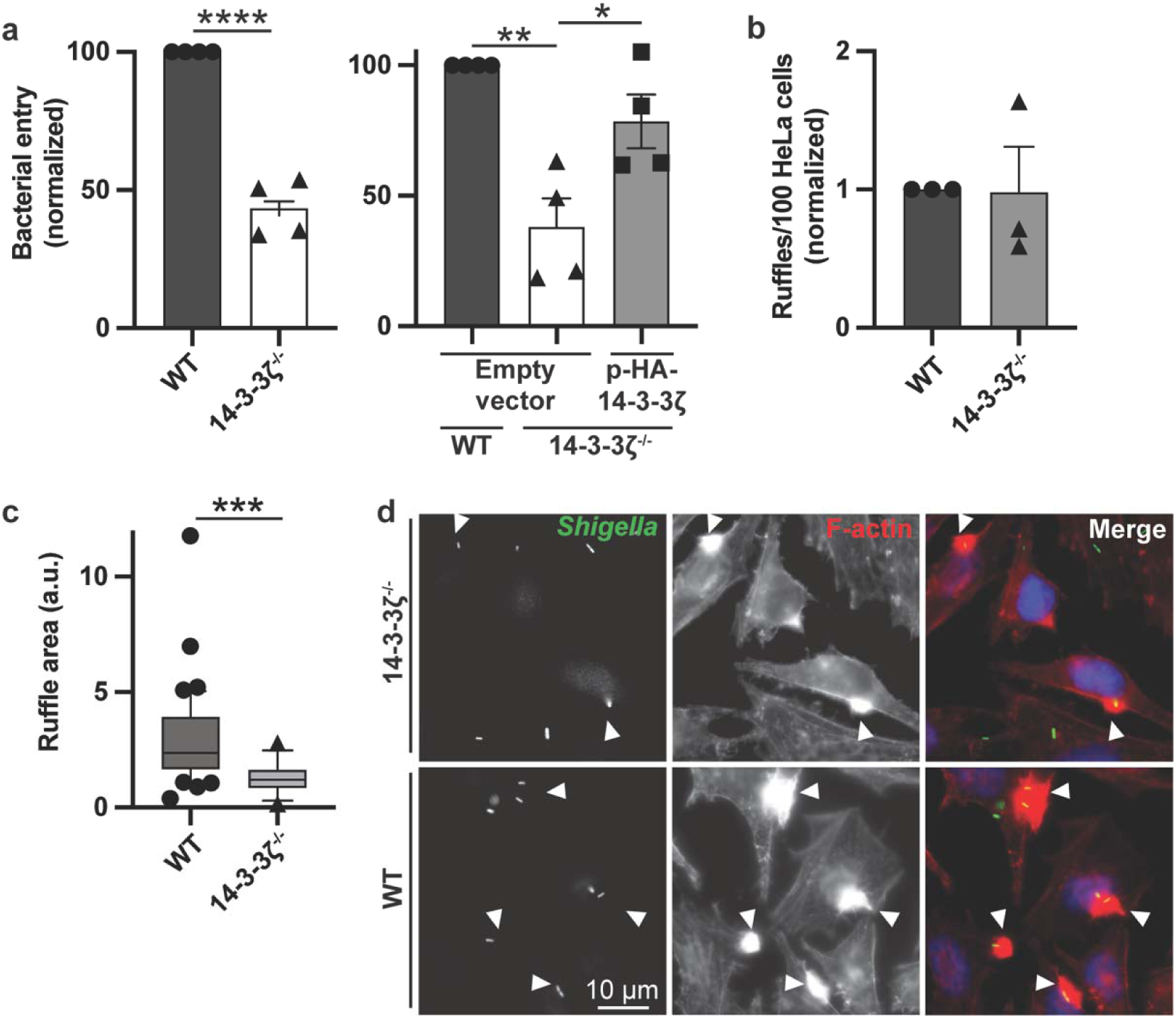
14-3-3ζ is required for efficient membrane ruffle formation and *S. flexneri* entry into cells. **a,** Reduction of *S. flexneri* entry into HeLa cells lacking 14-3-3ζ, which is rescued by complementation with 14-3-3ζ. Left graph, entry efficiency defined as the number of plaques formed in cell monolayers upon low MOI infection (see Methods), normalized to WT *S. flexneri*. Right graph, rescue of entry in 14-3-3ζ^-/-^ HeLa cells with complementation *in trans* by transfection of HA-tagged 14-3-3ζ. N = 4 independent biological replicates. **b,** 14-3-3ζ^-/-^ HeLa cells support bacterial-induced membrane formation. Number of ruffles per 100 HeLa cells, normalized to WT cells. N = 3 independent biological replicates. **c,** Reduction in area of plasma membrane ruffles induced during *S. flexneri* infection of cells lacking 14-3-3. Box plot of median value and interquartile range (IQR)1-IQR3 (25^th-^75^th^ percentile range of values). Whiskers represent 10^th^-90^th^ percentile of ruffles quantified. Individually plotted data points fall outside of percentile whiskers. Circles, HeLa WT; triangles HeLa 14-3-3ζ^-/-^. **d,** Visualization of reduction in size of ruffles during *S. flexneri* infection of cells lacking 14-3-3ζ. Fluorescence microscopy of infected 14-3-3ζ^-/-^ (top) or WT (bottom) HeLa cells, labeled for bacteria (green), polymerized actin (F-actin, phalloidin, red), and DNA (Hoechst, blue). Arrowheads, plasma membrane ruffles associated with bacteria. Mean ± SEM (**a**). *, p<0.05; **, p<0.01; ***, p<0.001; ****, p<0.0001.

### 14-3-3ζ functions as a molecular scaffold to support formation of bacterial-induced membrane ruffles

*S. flexneri* entry occurs within seconds to minutes of cell contact^13^. Membrane-inserted T3SS translocases interact with host intermediate filaments^13^, enabling delivery of effectors into the cytoplasm that induce membrane ruffle formation and bacterial uptake (Fig. 1b). To identify the step at which 14-3-3ζ contributes to entry, we tested the impact of the absence of 14-3-3ζ on specific steps in bacterial entry. In the absence of 14-3-3ζ, *S. flexneri* efficiently delivered and assembled functional, membrane-embedded translocons, indicating that the observed 14-3-3ζ-dependent defects in entry were not due to translocase instability or loss of function of membrane-embedded translocons (Extended Data Fig. 3d,e). However, the formation of membrane ruffles was defective; whereas membrane ruffles were present at infection foci in HeLa 14-3-3ζ^-/-^ monolayers (Fig. 2b), those ruffles that formed were significantly smaller than the ruffles that formed at infection foci in WT HeLa monolayers (Fig. 2c,d).

To test whether the scaffolding function of 14-3-3ζ contributes to *S. flexneri* membrane ruffle formation, we generated 14-3-3ζ mutants defective in dimerization^26^, vimentin binding^27^, and binding to phosphorylated proteins^28^ (Table 1). Complementation of HeLa 14-3-3ζ^-/-^ cells with HA-14-3-3ζ (WT) rescued the size of membrane ruffles to that observed in WT HeLa cells, whereas complementation with none of the 14-3-3ζ mutants rescued the defect (Fig. 3, Extended Data Fig. 2c,d). These data indicate that efficient membrane ruffle formation during *S. flexneri* entry requires dimerization and protein interaction capabilities of 14-3-3ζ, consistent with 14-3-3ζ functioning as a molecular scaffold for *S. flexneri* entry.

**Table 1.** 14-3-3ζ mutants.

| **Native sequence** | **Mutation** | **Mutant designation** | **Loss of function** |
| --- | --- | --- | --- |
| L<sub>12</sub>AE | Q<sub>12</sub>QR | Monomer | Dimerization<sup>26</sup> |
| E<sub>5</sub>, L<sub>12</sub>AE, Y<sub>82</sub>REKIE | K<sub>5</sub>, Q<sub>12</sub>QR,<br>Q<sub>82</sub>RENIQ | Vimentin | Vimentin binding<sup>27</sup> |
| K<sub>49</sub>, R<sub>56</sub>, R<sub>127</sub>, Y<sub>128</sub> | A<sub>49</sub>, A<sub>56</sub>, A<sub>127</sub>, A<sub>128</sub> | Binding pocket | Protein-protein interactions via phospho-binding pocket<sup>28</sup> |

**Fig. 3.**
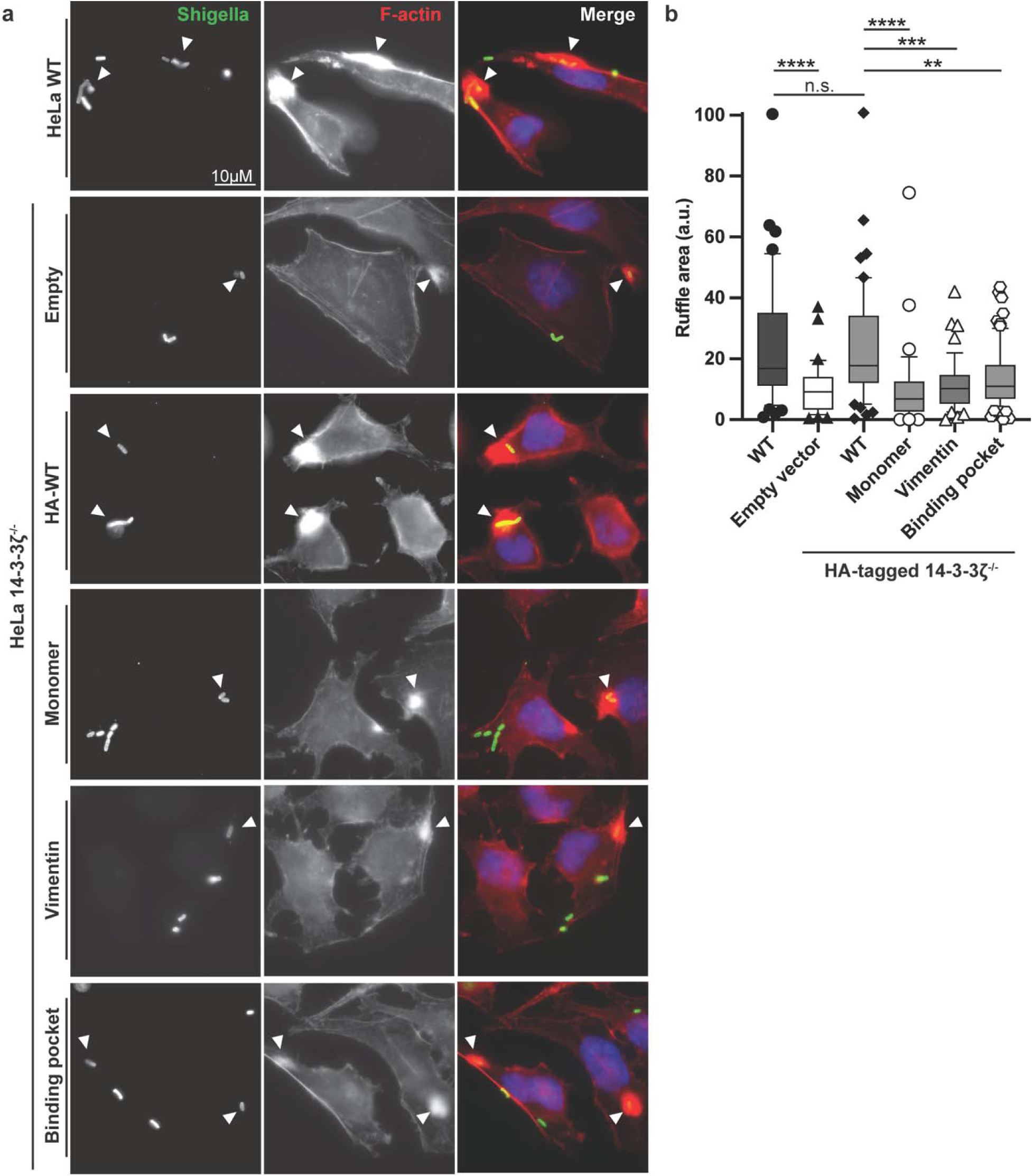
Derivatives of 14-3-3ζ with known functional defects support defective *S. flexneri*-induced membrane ruffles. **a,** Membrane ruffles induced during *S. flexneri* infection of HeLa cells expressing 14-3-3ζ derivatives defective in 14-3-3ζ dimerization (Monomer), in 14-3-3ζ interaction with vimentin (Vimentin), or in 14-3-3ζ binding to phosphorylated ligands (Binding pocket). Fluorescence microscopy of infection of HA-tagged WT or mutant 14-3-3ζ derivatives transfected into 14-3-3ζ^-/-^ HeLa cells. Bacteria (green), polymerized actin (phalloidin, red), and DNA (Hoechst, blue). **b,** Functionally defective 14-3-3ζ derivatives are associated with reductions in area of plasma membrane ruffles induced by *S. flexneri* infection of cells. WT HeLa cells, filled circles. HeLa 14-3-3ζ^-/-^ cells complemented with each of the following HA-tagged 14-3-3ζ derivatives: empty vector, filled triangles; WT 14-3-3ζ, filled diamonds; dimerization-defective 14-3-3ζ derivative, open circles; 14-3-3ζ derivative defective in vimentin binding, open triangles; 14-3-3ζ derivative defective in phospho-binding pocket, open hexagons. Whiskers represent 10^th^-90^th^ percentile of ruffles quantified. Individually plotted data points fall outside of percentile whiskers.

### Host proteins that depend on 14-3-3**ζ** for recruitment to translocons

To identify proteins that potentially function with 14-3-3ζ in *S. flexneri* entry, we compared VLP proteomes from WT HeLa monolayers to those of HeLa 14-3-3ζ^-/-^ monolayers, each infected with *S. flexneri* expressing functionally WT translocons, *S. flexneri* expressing functionally stalled translocons, or *S. flexneri* Δ*ipaC* (which do not form translocons due to lack of the translocase IpaC) (Fig. 4a, Supplementary Table 1). During infection with *S. flexneri* expressing WT and stalled translocons, CAP2 abundance was significantly enriched in VLPs derived from WT HeLa monolayers compared with those derived from HeLa 14-3-3ζ^-/-^ monolayers, indicating that CAP2 accumulation at the cytosolic face of translocons depends at least in part on 14-3-3ζ. This enrichment was not observed in VLPs from uninfected or *S. flexneri* Δ*ipaC*-infected monolayers (Fig. 4b), indicating that CAP2 abundance in the cell cortex is dependent on the presence of assembled translocons.

**Fig. 4.**
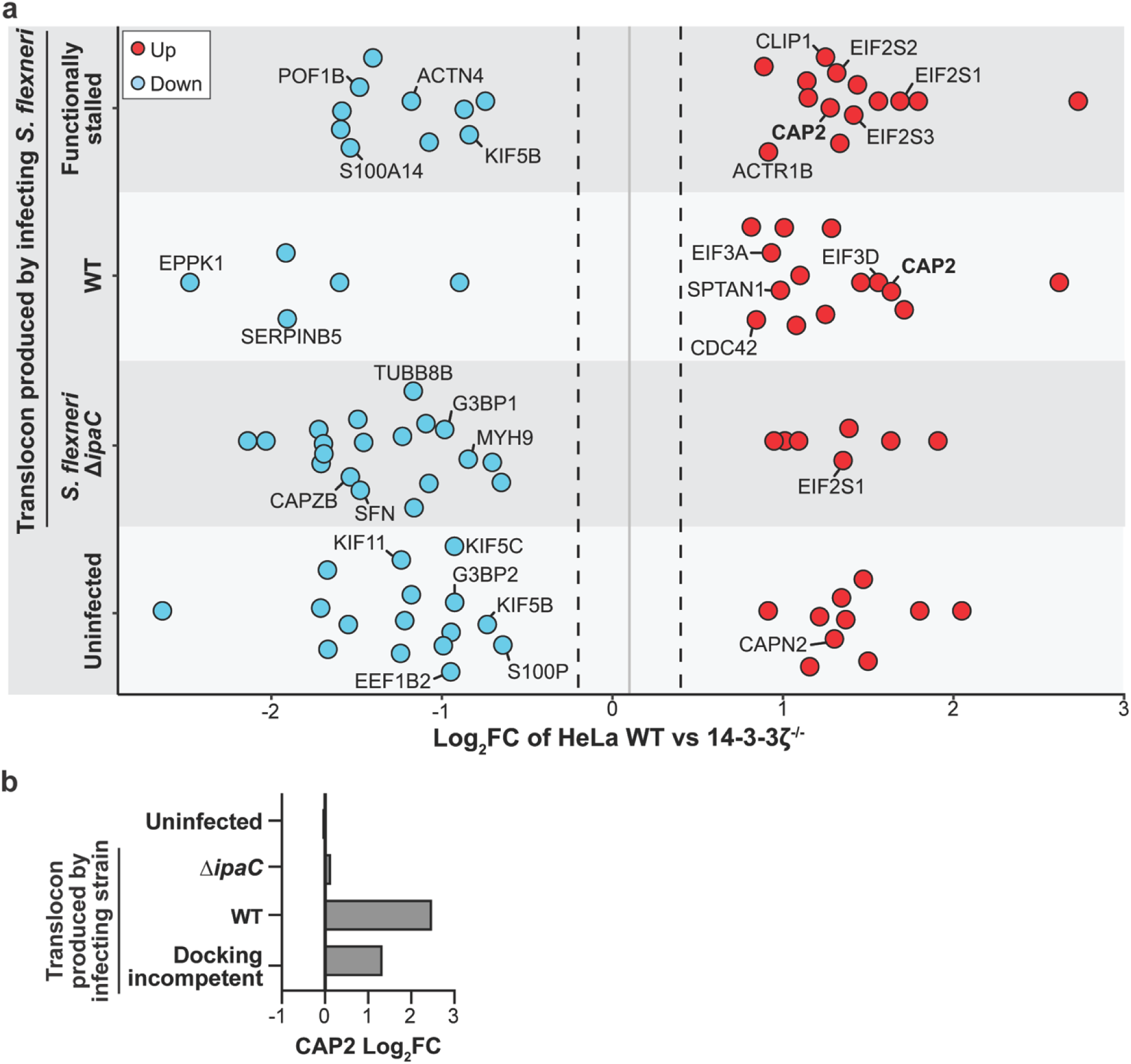
Proteomic comparison of VLPs from 14-3-3ζ^-/-^ cells to WT cells during *S. flexneri* entry. **a,** Enrichment of cellular proteins in VLPs from WT HeLa cells relative to HeLa 14-3-3ζ^-/-^ cells, infected with *S. flexneri* producing functional or stalled type 3 translocons or not producing intact translocons due to lack of the translocase IpaC. Mass spectrometry analysis of VLPs. Shown are significant (p ≤ 0.05) Log_2_FC in protein abundance. Labeled are cytoskeletal proteins and proteins known to participate in *Shigella* pathogenesis. Bold, CAP2. **b,** Increased abundance of CAP2 in VLPs from WT cells relative to VLPs from 14-3-3ζ^-/-^ cells during infection with WT or docking incompetent *S. flexneri*, each of which inserts translocons efficiently into the plasma membrane. Shown is Log_2_FC in CAP2 abundance.

### CAP2 localizes to membrane-engaged *S. flexneri* in a membrane ruffle-dependent manner

CAP2 is one of several proteins that control the cellular cytoskeletal actin network^29^. During *S. flexneri* infection, CAP2 colocalized with actin foci in bacteria-associated membrane ruffles in a 14-3-3ζ-independent manner (Pearson’s R = 0.79 ± 0.21 and 0.66 ± 0.19 in WT and 14-3-3ζ^-/-^ cells, respectively) (Fig. 5a,b). However, VLP proteomics (Fig. 4a), which may be more sensitive than immunofluorescence, showed that CAP2 accumulation is dependent in part upon the presence of 14-3-3ζ. In uninfected cells, CAP2 was distributed throughout the cytosol, only weakly colocalized with cortical actin, indicating that CAP2 enrichment in the actin cortex depends on factors involved in bacterial entry. These results indicate that bacterial engagement with the plasma membrane triggers CAP2 localization to bacterial-induced membrane ruffles in a manner that is likely enhanced by 14-3-3ζ.

**Fig. 5.**
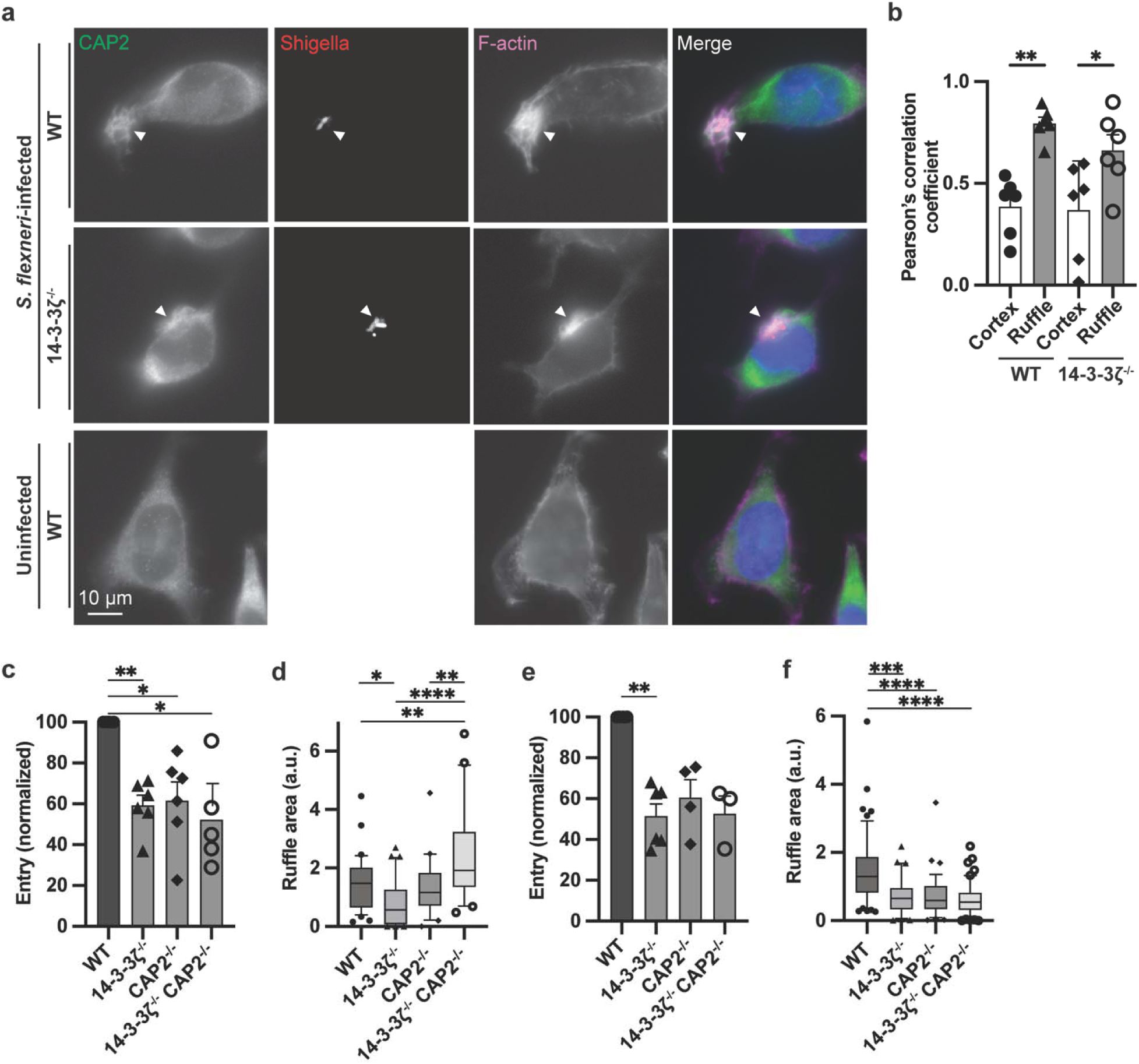
CAP2 and 14-3-3ζ act in the same pathway to promote membrane ruffle formation during T3SS pathogen infection. **a,** Fluorescence microscopy localization of CAP2 and polymerized actin with and without *S. flexneri* infection. CAP2 (green), bacteria (red), polymerized actin (far red). **b,** Peason’s correlation coefficient of CAP2 colocalization with polymerized actin in the indicated conditions. **c,** Absence of 14-3-3ζ and/or CAP2 lead to defects in *S. flexneri* entry. Entry is defined as the number of bacteria recovered from the monolayer at 90 mins of infection in the presence of gentamicin (see Methods). Normalized to WT HeLa. N = 5-6 independent biological replicates. **d,** Area of membrane ruffles induced during *S. flexneri* infection. Box plot representing median values, IQR1, and IQR3. Whiskers represent 10^th^-90^th^ percentile of ruffles quantified. Individually plotted data points fall outside of percentile whiskers. **e,** Absence of 14-3-3ζ and/or CAP2 leads to defects in *S.* Typhimurium entry. N = 3-6 independent biological replicates. **f,** Area of membrane ruffles induced during *S.* Typhimurium infection. Defined as in panel **d**. HeLa cells (**c-f**): WT, filled circles; 14-3-3ζ^-/-^, filled triangles; CAP2^-/-^, filled diamonds; 14-3-3ζ^-/-^ CAP2^-/-^, open circles.

### T3SS-induced membrane ruffle formation is coordinately regulated by 14-3-3**ζ** and CAP2

Using HeLa CAP2^-/-^ and 14-3-3ζ^-/-^ CAP2^-/-^ cells, we tested the effect of CAP2 on bacterial entry. Bacterial entry into CAP2^-/-^ and 14-3-3ζ^-/-^ CAP2^-/-^ monolayers was reduced by 39% ± 9 and 48% ± 11, respectively, compared with HeLa WT monolayers, similar to the entry defect observed for 14-3-3ζ^-/-^ monolayers (41% ± 5) (Fig. 5c). This result indicates that both 14-3-3ζ and CAP2 are required for efficient *S. flexneri* entry. The finding that the entry defect observed during infection of 14-3-3ζ^-/-^ CAP2^-/-^ cells was not additive of the defects that occurred during infection of 14-3-3ζ^-/-^ and CAP2^-/-^ cells suggests that CAP2 and 14-3-3ζ act in the same signaling pathway during T3SS pathogen entry.

Membrane ruffles formed during *S. flexneri* entry of HeLa 14-3-3ζ^-/-^ monolayers were significantly smaller than those formed in WT HeLa monolayers (Fig. 2d). While absence of CAP2 alone did not lead to significant changes in membrane ruffle size, bacterial-associated membrane ruffles in HeLa 14-3-3ζ^-/-^ CAP2^-/-^ monolayers were significantly larger than HeLa WT, 14-3-3ζ^-/-^, and CAP2^-/-^ monolayers (Fig. 5d). Altogether these results suggest that both 14-3-3ζ and CAP2 are involved in regulating membrane ruffle formation during bacterial uptake and support a model where CAP2 activity in membrane ruffle formation is independent of 14-3-3ζ but that CAP2 accumulation in membrane ruffles is enhanced by the scaffolding activity of 14-3-3ζ dimers.

To determine whether our observations are relevant to other T3SS pathogens, we tested the requirement for 14-3-3ζ and CAP2 during entry of *Salmonella enterica* Typhimurium SL1344 (*S*. Typhimurium). As with entry of *S. flexneri*, the absence of 14-3-3ζ and CAP2, both individually and together, reduced *S*. Typhimurium entry by 49% ± 6, 39% ± 9, and 47% ± 9, respectively, compared with WT HeLa monolayers (Fig. 5e). These data indicate that 14-3-3ζ and CAP2 are required for efficient bacterial entry not only for *S. flexneri* but also for another T3SS pathogen. Of note, where *S. flexneri*-induced membrane ruffles were significantly larger in HeLa 14-3-3ζ^-/-^ CAP2^-/-^ monolayers than in WT HeLa monolayers, *S*. Typhimurium-induced membrane ruffles were significantly smaller in 14-3-3ζ^-/-^, CAP2^-/-^, and 14-3-3ζ^-/-^ CAP2^-/-^ monolayers than in WT HeLa monolayers (Fig. 5f).

## Discussion

Among the challenges in identifying novel intracellular protein-protein interactions relevant to bacterial infection of eukaryotic cells are the capture of transient or dynamic interactions and discrimination of target complexes from the sheer volume of the cellular proteome^30,31^. A tool that combats these challenges is Virotrap, a method that fuses the protein target of interest to HIV Gag, which coopts the HIV budding mechanism, encapsulating both stable and transient target protein complexes in extracellular VLPs that can be immunoprecipitated for proteomic analyses^18,19^. Use of this approach to identify host factors required for bacterial pathogenesis has been limited^32^. Whereas these previously published Virotrap methods utilized FLAG-VSVG for VLP immunoprecipitation, here, to identify host proteins specifically recruited to T3SS translocons during bacterial entry, we adapted Virotrap to capture *S. flexneri* IpaB-FLAG-containing VLPs following native delivery of bacterial T3SS translocons to HeLa cells that express Gag. Gag-expressing HeLa cells support translocon insertion and production of translocon-containing VLPs, and the resulting VLPs yielded new insights into host proteins that function in translocon-induced bacterial uptake, establishing that this adapted Virotrap method is feasible and productive in the study of interactions between bacterial transmembrane proteins and host proteins.

The plasma membrane-embedded T3SS translocon is a dynamic structure that undergoes conformational changes upon interaction with the host cytoskeleton, leading to membrane ruffling and bacterial internalization by micropinocytosis^10,13^. Several host factors that contribute to these interactions have been described^10,13,14,16,17,33^, yet the process remains incompletely understood. By capturing in VLPs host proteins present at the cytosolic face of *S. flexneri* translocons, we identified additional host factors required for *S. flexneri* manipulation of host membranes and the cortical cytoskeleton (Fig. 1a)^4,34,35^. To maximize proteomic recovery of biologically relevant host proteins recruited to the translocon, we leveraged our prior identification of a translocon point mutant that forms functionally stalled translocons, thereby locking the translocon in a docking incompetent state (Fig. 1b)^13^.

Among the proteins we found enriched in the proteome of VLPs containing functionally stalled translocons, and which was required for efficient bacterial entry and membrane ruffle formation, was the 14-3-3 protein 14-3-3ζ. 14-3-3 proteins are small, soluble, intracellular proteins that participate in signaling pathways via binding to phosphoserine and/or phosphothreonine when in monomer or dimer form^21,22,36,37^. 14-3-3ζ is required for activation of *P. aeruginosa* ExoS during infection^23^, by mechanisms unrelated to its role in *S. flexneri* and *S.* Typhimurium entry. Whereas 14-3-3 isoforms have been thought to be functionally redundant, individual 14-3-3 isoforms display differential tissue specificity and subcellular localization^38^. All isoforms are present in HeLa cells (Extended Data Fig. 2b), and we found a specific requirement for 14-3-3ζ in *S. flexneri* and *S*. Typhimurium entry. Thus, with this study, we attribute a novel function to 14-3-3ζ that is non-redundant with other isoforms.

14-3-3 proteins are maintained at steady levels in the cytoskeletal network via binding to phosphorylated vimentin^27^, an intermediate filament. *S. flexneri* T3SS translocons also bind intermediate filaments^13^, consistent with engagement of the same cytoskeletal compartment. Our finding that 14-3-3ζ was significantly enriched in VLPs and was required for efficient membrane ruffle formation during *S. flexneri* entry (Fig. 2) indicates that 14-3-3ζ contributes to bacterial entry through modulation of the membrane ruffle cytoskeletal architecture.

Among their roles in signaling pathways, 14-3-3 proteins act as molecular scaffolds that drive protein-protein interactions and signaling mediated by these interactions^21^. Several mutations in 14-3-3, including defects in dimerization and vimentin binding, and abrogation of the phospho-binding pocket, are known to cause specific functional defects (Table 1)^28^. Complementation of HeLa 14-3-3ζ^-/-^ with each mutant construct did not rescue *S. flexneri*-induced membrane ruffle morphology (Fig. 3, Extended Data Fig. 2c,d), indicating that 14-3-3ζ functions as a molecular scaffold in membrane ruffles during infection and that 14-3-3ζ interactions with other cytoskeletal components are critical for membrane ruffle formation.

To identify proteins that interact with 14-3-3ζ during bacterial entry, we compared the proteomes of translocon-containing VLPs from HeLa WT cells against VLPs from 14-3-3ζ^-/-^ cells (Fig. 4, Supplementary Table 2). Among the proteins we found to be enriched and required for efficient bacterial entry is CAP2, a capping protein that accelerates actin turnover by promoting release of G-actin from the ends of actin filaments^39,40^ and nucleotide exchange on G-actin^41,42^, which prepares the monomers for rapid incorporation into growing filaments. Cells lacking CAP2 exhibit increased membrane protrusions and focal adhesions and show defects in cell migration^29^. CAP2 is best understood in the context of cardiac disease, as CAP2 mutations and/or deletions are associated with dilated cardiomyopathy and dysfunction of the sarcomere^43,44^. A role for CAP2 in bacterial infection has not been previously reported.

Based on our findings that absence of 14-3-3ζ or CAP2 alone lead to bacterial entry defects, but that the dual absence of 14-3-3ζ and CAP2 did not result in an additive entry defect (Fig. 5c), we propose that 14-3-3ζ and CAP2 act in the same pathway during bacterial entry. Although we found by immunofluorescence microscopy that CAP2 localized to membrane ruffles even in the absence of 14-3-3ζ (Fig. 5a), the marked decrease in enrichment of CAP2 in VLPs from 14-3-3ζ^-/-^ cells (Fig. 4a) indicates that CAP2 recruitment to sites of *S. flexneri* entry is at least in part dependent on 14-3-3ζ, suggesting that 14-3-3ζ may act upstream of CAP2 during membrane ruffle formation. It is possible that in the absence of 14-3-3ζ, CAP2 is recruited into the active cortical actin network, and that 14-3-3ζ brings CAP2 closer to the plasma membrane, leading to improved incorporation into VLPs. Altogether these data are consistent with a model in which upon translocon insertion, 14-3-3ζ and CAP2 coordinately contribute to the formation of membrane ruffles capable of bacterial uptake.

We tested whether the roles of 14-3-3ζ and CAP2 in bacterial entry are conserved among T3SS translocon homologs using *S*. Typhimurium. As for *S. flexneri*, both 14-3-3ζ and CAP2 were necessary for efficient *S*. Typhimurium entry (Fig. 5c,e). In contrast, whereas only 14-3-3ζ modulated *S. flexneri* membrane ruffle size, *S*. Typhimurium infection of CAP2^-/-^ monolayers resulted in smaller membrane ruffles comparable to those produced in WT monolayers (Fig. 5d,f). Membrane ruffle expansion during *S*. Typhimurium infection requires vesicular delivery of plasma membrane reservoirs via Rab10 and LRRK2, without which the ruffles remain stunted and unable to support bacterial internalization^45,46^. Although it is not known whether Rab10 and/or LRRK2 are required during *S. flexneri* infection, these findings raise the possibility of multiple 14-3-3 functions during *S*. Typhimurium infection giving rise to infection phenotypes distinct from those observed during *S. flexneri* infection. 14-3-3 dimers negatively regulate LRRK2 activity^47^, yet whether this specific activity of 14-3-3 is altered during pathogen infection is unknown.

14-3-3ζ, either directly or indirectly, promotes recruitment of CAP2 to bacterial entry sites. 14-3-3ζ is known to promote lamellipodia and membrane ruffle formation through the cytoskeletal binding partner ARHGEF7 (Rho guanine nucleotide exchange factor 7, βPIX)^48^. Rac1, the small GTPase that activates lamellipodia formation, is targeted by ARHGEF7 and is activated by *Shigella* and *Salmonella* effectors in a manner that enhances entry^16,49^. In the presence of a dominant-negative derivative of 14-3-3ζ, instead of promoting lamellipodia, ARHGEF7 promotes filopodia^48^, suggesting that in the absence of 14-3-3ζ, activation of Rac1 by the pathogen may lead to misdirection of actin polymerization into filopodia instead of lamellipodia, which would be expected to hamper bacterial entry.

We propose a model in which CAP2 generates pools of G-actin for *Shigella*-directed (or *Salmonella*-directed) actin polymerization for membrane ruffle formation at entry sites. In the absence of CAP2, the pool of G-actin is reduced, leading to less effective bacterial direction of membrane ruffle generation. We found that *S. flexneri* formed significantly larger membrane ruffles during infection of HeLa monolayers lacking both 14-3-3ζ^-/-^ and CAP2^-/-^ (Fig. 5c). The absence of both might be expected to be associated with both a reduction in the CAP2-generated G-actin pool and hampered 14-3-3ζ-directed lamellipodia formation, which together with actin polymerization-promoting bacterial translocase IpaC^50^ and effectors IpaA^51^ and IpgB1^52^, would likely promote misdirected filopodia formation; our findings in the dual absence of 14-3-3ζ and CAP2 are consistent with this. Although larger, the ruffles present in the HeLa 14-3-3ζ^-/-^ CAP2^-/-^ monolayers did not support recovery of bacterial entry compared with monolayers lacking each protein individually (Fig. 5c), consistent with these large ruffles lacking the structural capacity for efficient *S. flexneri* internalization and highlighting that the precise coordination of bacterially-induced membrane ruffles is required for pathogenesis.

Altogether our data show that 14-3-3ζ and CAP2 work together to regulate membrane ruffle dynamics during *S. flexneri* infection. These data are consistent with a model in which CAP2 is recruited to sites of bacterial entry (in part by 14-3-3ζ), promoting bacterial effector-induced lamellipodia and associated membrane ruffles, which support bacterial entry. Our data indicate that T3SS pathogens co-opt 14-3-3ζ and CAP2-associated signaling pathways during host entry. In the context of infection, it is likely that these two events happen rapidly upon T3SS translocon insertion into host plasma membranes and the resulting cytoskeletal remodeling triggers 14-3-3ζ dimer scaffolding and CAP2 recruitment and activation to manipulate the cytoskeletal network within membrane ruffles towards bacterial uptake. These mechanisms are likely to be broadly relevant to T3SS pathogens and non T3SS pathogens that manipulate host cytoskeletal networks during host cell entry.

## Online Methods

### Bacterial Strains and Plasmids

The bacterial strains and plasmids used in this study are listed in Supplementary Table 3. Wildtype *S. flexneri* serotype 2a strain 2457T has been described^53^. All mutants are isogenic derivatives of 2457T. The Δ*ipaB* Δ*ipaC* mutant was generated using the Wanner method^54^. IpaB and IpaC derivatives were encoded by genes inserted into the plasmid pDSW206^55^, for which protein expression is driven by the ColE1 promoter and induced with 1mM isopropyl thio-β-galactoside (IPTG); these plasmids were introduced into *S. flexneri* Δ*ipaB* Δ*ipaC* and its derivatives, as appropriate. Unless otherwise noted, to enhance translocon delivery into cell membranes, all *S. flexneri* strains expressed the *E. coli* adhesion Afa-1^56^. The sequences of primers used in PCR and sequencing are available from the authors upon request.

### Cell Culture

HeLa (CCL2) cells were obtained from ATCC, cultured in Dulbecco’s Modified Eagles media (DMEM) supplemented with 0.45% glucose and 10% heat-inactivated fetal bovine serum (FBS), and grown at 37°C with 5% CO_2_ and humidity.

### Quantification of T3SS translocases in plasma membranes

Membrane isolation and detection of translocon proteins was performed as described previously^8,57^. Briefly, 3 x 10^5^ HeLa cells were seeded into each well of a 6-well plate. Two wells were used for each strain tested. *S. flexneri* cultures were diluted to an MOI of 100 in PBS with 1mM IPTG. Bacteria were centrifuged onto HeLa monolayers at 25°C for 10 minutes at 2,000 x *g*, and infected cells were incubated at 37°C in 5% CO2 with humidified air for 20 minutes. Infected cells were subsequently harvested, and plasma membrane-enriched fractions, which contains translocons, were isolated via sequential membrane fractionation with 0.2% saponin then 0.5% TritonX-100, as previously described^9,58^. Western blots were performed using rabbit anti-caveolin-1 (Sigma, C4490), rabbit anti-GroEL (Sigma, G6532), rabbit anti-IpaC (gift of R. Kaminski), and mouse anti-IpaB (gift of R. Kaminski).

### Bacterial effector translocation assay

Effector translocation was performed as previously described^10^. Briefly, 3 x 10^5^ HeLa cells were seeded into each well of a 6-well plate. Two wells were used for each strain tested. *S. flexneri* cultures expressing OspB-FLAG were diluted to an MOI of 100 in HBSS with 4% FBS and 1mM IPTG. Infection was carried out as described in Methods (Quantification of T3SS translocases in plasma membranes), and infected cells were incubated 37°C in 5% CO2 for 60 minutes before harvesting and lysing with RIPA buffer (50 mM Tris, pH 8 containing 150 mM NaCl, 1% Nonidet-P40, 0.1% SDS, 10 mM NaF, and EDTA-free protease inhibitor cocktail, Roche). Cellular debris and bacteria were separated by centrifugation and collected as the bacterial fraction, and the soluble fraction was collected as the cell cytosol. Abundance of OspB-FLAG delivered to the cytosol was determined by western blot using mouse anti-FLAG (Sigma, F1804), mouse anti-HIV1 p24 (Abcam, ab9071), mouse anti-β-actin-peroxidase (Sigma, A3854).

### Pore formation

Pore formation in nucleated cells was performed as described previously^59^. HeLa cells seeded in 96-well flat-bottom plates at 1.5×10^4^ per well, were preloaded with BCECF, AM [2’-7’-Bis-(2-Carboxyethyl)-5-(and-6)-Carboxyfluorescein, Acetoxymethyl Ester (Invitrogen, B1170)] and Hoechst (Invitrogen) in HBSS at 37°C with 5% CO_2_. The cells were infected with exponential phase bacteria at an MOI of 100 in RPMI. Entry was synchronized by centrifugation at 800x*g* for 10 minutes at 25°C, and the infection was continued at 37°C with 5% CO_2_ for 50 minutes. Fluorescent microscopy was used to image the total number of cells (blue filter) and BCECF-positive cells (green filter).

### Virotrap

10^7^ HeLa cells were transiently transfected with pMET7-GAG-GW and pcDNA3-FLAG-VSVG or pcDNA3-VSVG (this study) at a 9:1 ratio using FuGENE HD Transfection Reagent (Promega). pMET7-GAG-GW was a gift from Sven Eyckerman (Addgene plasmid # 194634; http://n2t.net/addgene:194634; RRID: Addgene_194634) and pcDNA3-FLAG-VSVG was a gift from Sven Eyckerman (Addgene plasmid # 80606; http://n2t.net/addgene:80606; RRID: Addgene_80606). DNA mixed with FuGENE HD at a 1:3.5 ratio in Opti-MEM (Gibco) was incubated at room temperature for 5 minutes, then added dropwise onto HeLa cells. pcDNA3-FLAG-VSVG was used in the DNA mix to generate VLPs for uninfected controls. pcDNA3-VSVG was used in the DNA mix to generate VLPs for infected samples. After 48 hours, transfected monolayers were washed with PBS and infected with *S. flexneri* expressing IpaB-FLAG at MOI 100. After 30 minutes of infection, gentamicin, which kills extracellular but not intracellular bacteria, was added to final concentration of 25μg/ml, and infected cells were incubated for an additional 2 hours. Cell culture supernatants, which contain VLPs, were collected and processed as previously described^19^. Briefly, cell debris was removed from supernatants by centrifugation and passage through 0.45μm filters. Supernatants were then filter concentrated using Amicon ultra centrifugal filters, 100kDa MWCO (Millipore), loaded onto anti-FLAG M2 magnetic beads (Millipore), and incubated for 2 hours at 4°C with end-over-end rotation. VLPs captured with anti-FLAG beads were washed three times, resuspended in SDS sample buffer and boiled for 10 min. at 95°C. Samples were then loaded in 12.5% SDS PAGE gels, and gel bands were excised and submitted to the Taplin Mass Spectrometry Facility proteomics core for analyses.

### Mass spectrometry/Proteomics

At the Taplin Mass Spectrometry Facility, excised gel bands were cut into approximately 1 mm^3^ pieces. Gel pieces were then subjected to a modified in-gel trypsin digestion procedure^60^. Gel pieces were washed and dehydrated with acetonitrile for 10 min. followed by removal of acetonitrile. Pieces were then completely dried in a speed-vac. Rehydration of the gel pieces was with 50 mM ammonium bicarbonate solution containing 12.5 ng/µl modified sequencing-grade trypsin (Promega, Madison, WI) at 4°C. After 45 min., the excess trypsin solution was removed and replaced with 50 mM ammonium bicarbonate solution to just cover the gel pieces. Samples were then placed in a 37°C room overnight. Peptides were later extracted by removing the ammonium bicarbonate solution, followed by one wash with a solution containing 50% acetonitrile and 1% formic acid. The extracts were then dried in a speed-vac (∼1 hr). The samples were then stored at 4°C until analysis.

On the day of analysis, the samples were reconstituted in 5 - 10 µl of HPLC solvent A (2.5% acetonitrile, 0.1% formic acid). A nano-scale reverse-phase HPLC capillary column was created by packing 2.6 µm C18 spherical silica beads into a fused silica capillary (100 µm inner diameter x ∼30 cm length) with a flame-drawn tip^61^. After equilibrating the column each sample was loaded via a Thermo EASY-LC (Thermo Fisher Scientific, Waltham, MA). A gradient was formed, and peptides were eluted with increasing concentrations of solvent B (90% acetonitrile, 0.1% formic acid).

As peptides eluted, they were subjected to electrospray ionization and then entered into a Orbitrap Fusion Lumos mass spectrometer (Thermo Fisher Scientific, Waltham, MA). Peptides were detected, isolated, and fragmented to produce a tandem mass spectrum of specific fragment ions for each peptide. Peptide sequences (and hence protein identity) were determined by matching protein databases with the acquired fragmentation pattern by the software program, Sequest (Thermo Fisher Scientific, Waltham, MA)^62^. All databases include a reversed version of all the sequences and the data was filtered to between a one and two percent peptide false discovery rate.

### Generation of CRISPR knockouts

To generate the HeLa 14-3-3ζ^-/-^ cell line, guide RNA sequence (“CD.Cas9 HRNB5585 AA” AGGAGATTACTACCGTTACT/GGGATTGTCGATCAGTCACA) was subcloned into a lentiviral construct, and HeLa cells were transduced with lentivirus packaged with CRISPRMAX.

To generate the HeLa CAP2^-/-^ cell line, RNP complexes were assembled with pooled guide RNA sequences (“guide 1” UGACUGCUUACAGAAUGACG, “guide 2” CUUUCAGAGAGAGAAACCGG, “guide 3” ACAGCUAUCCAUCCAAGGGC) and Cas9 at a ratio of 3:1 in Resuspension buffer R. 2 × 10^5^ HeLa cells were pelleted, resuspended in Resuspension Buffer R, added to the assembled RNP complex or scrambled control. HeLa WT and HeLa 14-3-3ζ^-/-^ were electroporated using the Neon Transfection System (Invitrogen) according to the manufacturer’s instructions. Using a 10 μL Neon tip, cells were pulsed once at 1005 V and 35 ms pulse width. Electroporated cells were immediately resuspended in 2.5 mL of DMEM with 10% FBS and placed into wells of a 6-well plate. Cells were incubated for 72 hours at 37°C before analysis with the Inference for CRISPR edits tool (Synthego) and limiting dilution for selection of a clonal homozygous CAP2 deletion lineage.

Levels of 14-3-3ζ transcripts were determined by quantitative reverse transcriptase polymerase chain reaction (qRT-PCR). Transcripts were prepared using RNeasy Micro kit (Qiagen), cDNA was generated using the SuperScript IV VILO kit (Thermo Fisher Scientific), and PCR was performed with SsoFast EvaGreen Supermix kit (Bio-Rad) for 40 cycles. Primers used in qRT-PCR were YWHAZ F: TTCTTGATCCCCAATGCTTC and YWHAZ R: AGTTAAGGGCCAGACCCAGT. The housekeeping genes used for reference were MAP3K14 (F: ACAACGAGGGTGTCCTGCTC, and R: TCAGCTCCTCTGCCCGAAA) and HPRT (F: CCTGGCGTCGTGATTAGTGAT, and R: AGACGTTCAGTCCTGTCCATAA).

14-3-3ζ and CAP2 proteins were detected with rabbit anti-14-3-3ζ (LSBio #LS-C353025) and rabbit anti-CAP2 antibodies (Proteintech #15865-1-AP), with β-actin as a loading control. Other 14-3-3 isoforms were detected with rabbit anti-YWHAB (Invitrogen, MA5-32419), rabbit anti-YWHAE (Abcam, ab92311), rabbit anti-YWHAG (Invitrogen, PIMA535769), rabbit anti-YWHAH (Abcam, ab206292), rabbit anti-SFN (Abcam, ab193667), and rabbit anti-YWHAQ (Abcam, ab124909). HA-14-3-3ζ was detected with mouse anti-HA (BioLegend, 901501).

### Quantification of adherent and intracellular bacteria

HeLa cells were seeded at 10^5^ cells per well in 24-well plates the day prior to infection. *S. flexneri* not expressing *E. coli* Afa-1 was grown as described above and added to cell monolayers in DMEM at an MOI of 100. Bacteria were centrifuged onto cells for 10 min. at 2,000 rpm and incubated at 37°C with 5% CO2 for 30 min. Bacteria that did not invade the cells were removed by washing with phosphate-buffered saline (PBS). To quantify the number of adherent bacteria, infected monolayers were then treated with 0.25% trypsin and lysed in PBS containing 0.5% Triton X-100. Lysates containing the bacteria were plated on agar plates to quantify the number of adherent bacteria. To quantify the number of intracellular bacteria, infected HeLa cells were cultured for an additional hour at 37°C with fresh DMEM containing 25 μg/ml gentamicin. Infected monolayers were then washed with PBS, treated with trypsin, and lysed in PBS containing 0.5% Triton X-100 to release the intracellular bacteria. Lysates containing the bacteria were plated on agar plates to quantify the number of intracellular bacteria.

### Quantification of bacterial spread

The plaque assay was performed by seeding 5.5 × 10L cells per well in a 6-well plate, the day prior to infection, followed by infection with WT *S. flexneri* not expressing *E. coli* Afa-1 at a multiplicity of infection (MOI) of 0.001. The plates were centrifuged at 2,000 rpm for 10 min. at room temperature to promote infection and subsequently incubated at 37°C for 1.5 hours. Following incubation, the media was removed, and an agarose overlay containing DMEM, 10% fetal bovine serum (FBS), 25 µg/mL gentamicin, and 1% agarose was added to each well. The overlay was allowed to solidify at room temperature before incubating the plates at 37°C for 72 hours to allow plaque formation. For visualization, a neutral red agarose overlay was applied, consisting of DMEM, 10% FBS, 25 µg/mL gentamicin, 1.4% agarose, neutral red solution, and 6N HCl. Plates were incubated for an additional 6 hours before plaques were scanned. The plaques were subsequently analyzed by quantifying the number and area of each plaque to assess infection efficiency and spread.

### Immunofluorescence microscopy of membrane ruffles, CAP2, and *S. flexneri*

HeLa cells were seeded onto glass coverslips at 3×10^5^ cells per well of a 6-well plate. After 24 hours, HeLa monolayers were infected with *S. flexneri* at an MOI of 2 by centrifugation at 800x*g* for 10 min. at room temperature, followed by an additional 10 min. at 37°C with 5% CO2. Infected cells were washed 3x with PBS then fixed with 4% paraformaldehyde for 20 min. at room temperature. HeLa membranes were permeabilized with 0.5% TritonX-100 for 7 min. at room temperature. DNA was stained with Hoechst, actin was stained with Alexa Fluor 568 conjugated to phalloidin (Invitrogen, A12380) or Alexa Fluor 647 conjugated to phalloidin (Invitrogen, A22287), CAP2 was stained with rabbit anti-CAP2 and an Alexa Fluor 488 secondary antibody (Invitrogen, A-11034), and *S. flexneri* was stained with rabbit anti-*Shigella* FITC (Virostat, 0903) conjugated to FITC. Coverslips were mounted onto glass slides with ProLong Diamond (Invitrogen) and images were collected by epifluorescence microscopy.

### Microscopy and image analysis

Images were collected using a Nikon TE-300 microscope equipped with Q-Imaging Exi Blue Cameras (Q-imaging), Chroma Filters, and IVision Software (BioVision Technologies). Unless noted otherwise, images were randomly collected across a coverslip. Single channel images were pseudo-colored and assembled in ImageJ or Illustrator (Adobe).

Signal from Western blots was captured by film or with an Azure 300 Chemiluminescent Imager (Azure Biosystems). Film was digitized using an Epson Perfection 4990 photo scanner, and the density of bands was determined using ImageJ (National Institutes of Health).

### Statistical analysis

Except where specifically noted, all data are from three independent experiments and the means ± standard errors of the means (SEM) are presented. Dots within graphs represent independent experimental replicates. The means between groups were compared by a one-way analysis of variance (ANOVA) using GraphPad Prism 8 (GraphPad Software, Inc.). Analysis of VLP proteomes was done by the Joslin Diabetes Center Bioinformatics and Biostatistics Core.

## Supporting information

Compiled supplementary data and tables excluding Supplementary table 2

## Acknowledgements

We thank Goldberg laboratory members for critical reading of the manuscript. We thank Robert Kaminski for reagents. We thank Ross Tomaino from the Taplin Mass Spectrometry Facility, Cell Biology Department, Harvard Medical School for mass spectrometry support. We thank Jonathan M. Dreyfuss and Hui Pan from the Joslin Diabetes Center’s Bioinformatics & Biostatistics Core for proteomics data analysis and support.

This work was funded by NIH grant R01 AI081724 (to M.B.G.) and a Massachusetts General Hospital Executive Committee on Research Fund for Medical Discovery Fundamental Research Fellowship Award (to M.L.G.M.).

The funders had no role in study design, data collection and interpretation, or the decision to submit the work for publication. The authors have no financial or other relationships that are relevant to the study.

Study conceptualization was by P.C. and M.B.G.; investigation by P.C., M.L.G.M., M.S.F., L.A.T.P., K.T.E., and V.K.; methodology and formal analysis by P.C. and M.L.G.M.; writing of the original draft by P.C.; and review and editing by M.L.G.M, M.S.F., L.A.T.P., K.T.E., V.K., and M.B.G.

