## Supplementary material for "Virus-like particle capture reveals coordination of actin remodeling during Shigella flexneri entry by host proteins": Compiled supplementary data and tables excluding Supplementary table 2

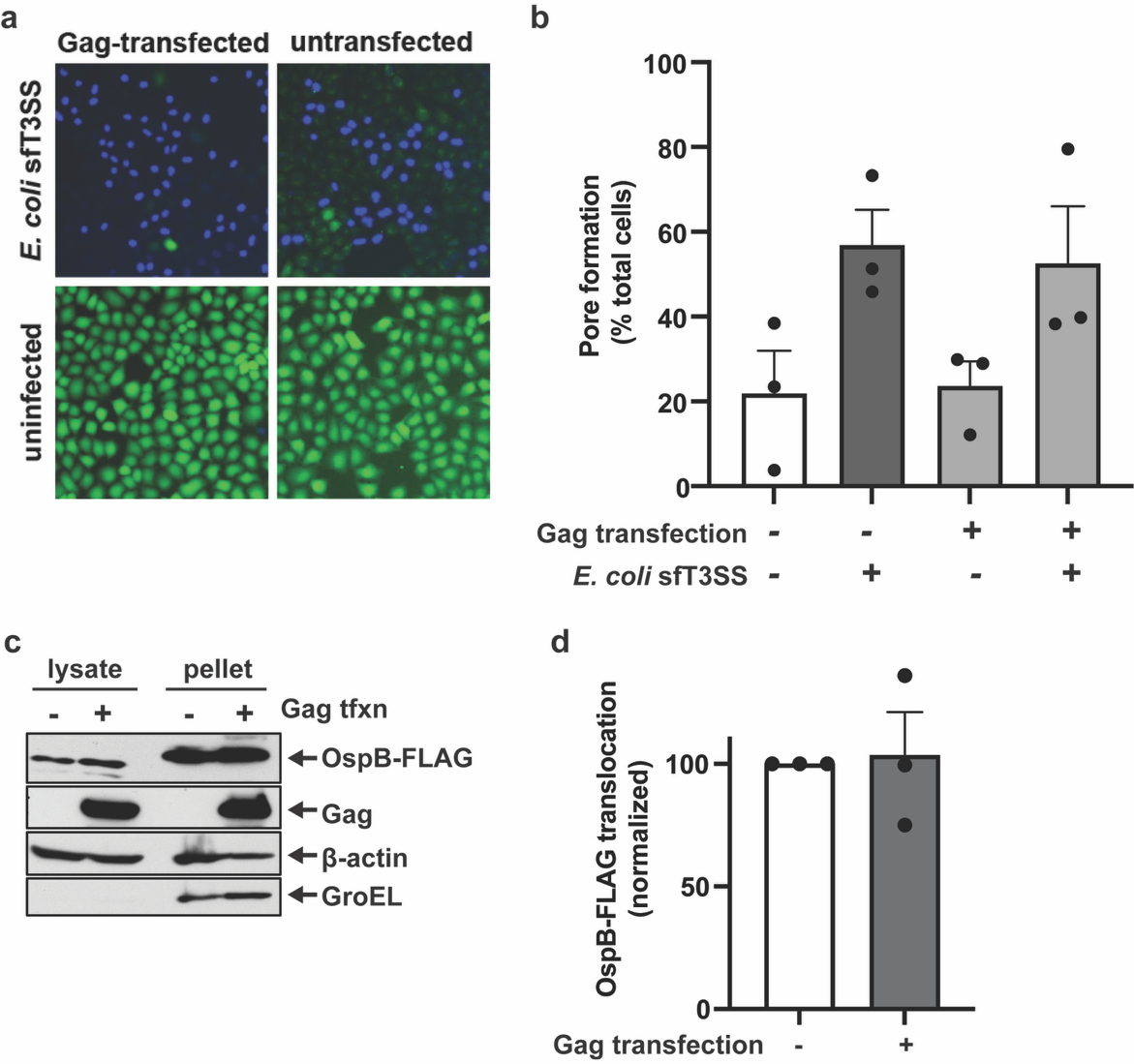

**Supplementary Fig. 1. Adaptation of Virotrap to generate viral-like particles (VLPs) that capture *S. flexneri* invasion.** **a-b,** Formation of pores by the *S. flexneri* type 3 translocon in the plasma membranes of HeLa cells is unaffected by transfection with Gag. Release of pre-loaded BCECF dye (green) from HeLa cells transfected or not with Gag upon infection with *E. coli* DH10B that deliver into plasma membranes the *S. flexneri* type 3 translocon (pSfT3SS)^63^. DNA (blue, Hoechst) (**a**). Quantification of pore formation as the percentage of cells having lost green fluorescence. N = 3 independent biological replicates (**b**). **c-d,** Effector translocation via type 3 secretion during *S. flexneri* infection is unaffected by transfection with Gag. Presence of effector OspB-FLAG in soluble lysates of infected cells transfected or not transfected with Gag (lanes 1-2, top blot); intact bacteria segregate with pellets, as demonstrated by restriction of bacterial cytoplasmic protein GroEL to the pellets (lanes 3-4, bottom blot). Western blot analysis with antibodies to FLAG, Gag, β-actin, or GroEL (**c**). Band densitometry of OspB-FLAG in soluble fractions. Normalized to no transfection of Gag. N = 3 independent biological replicates (**d**). Mean ± SEM (**b, d**).

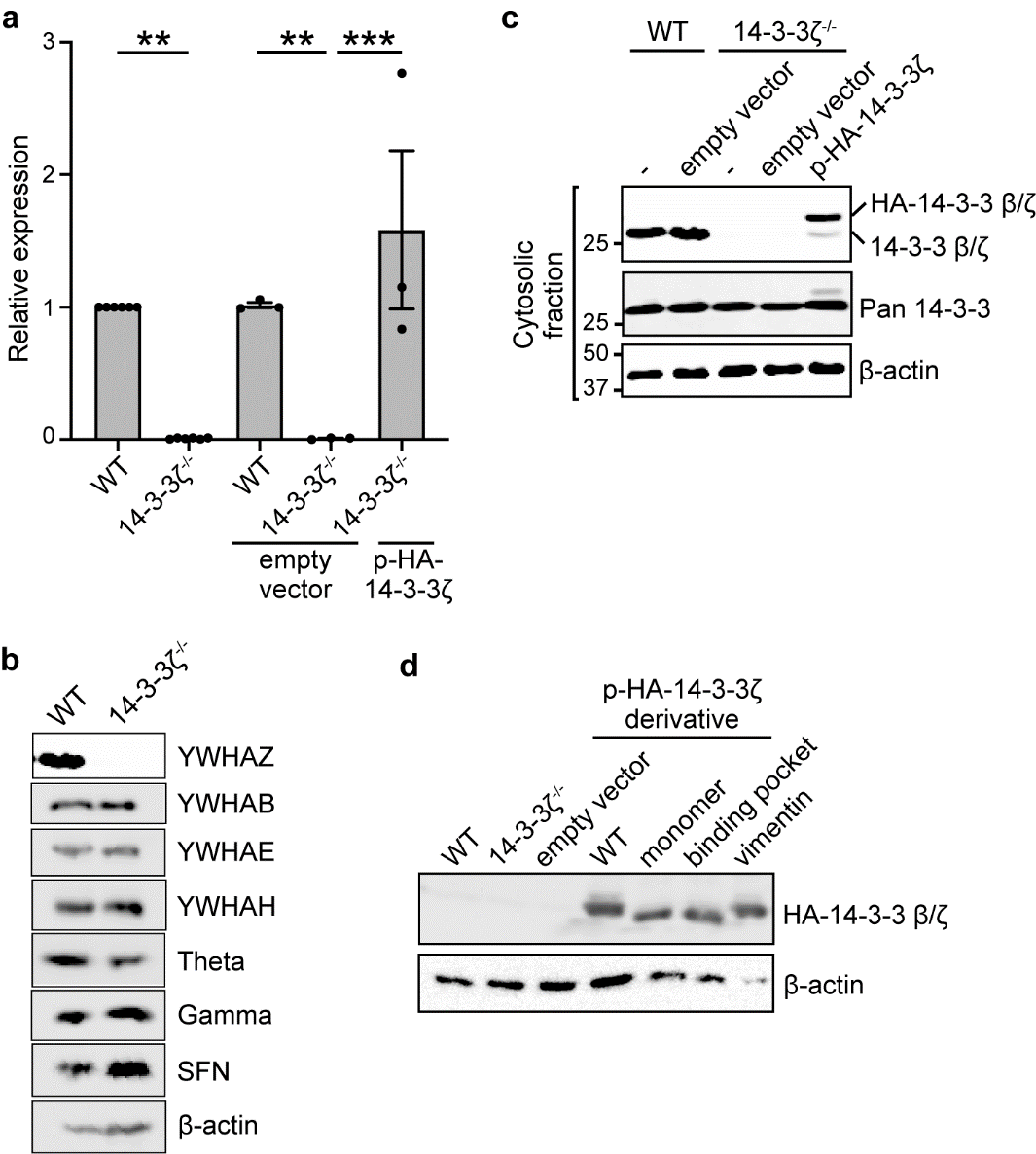

**Supplementary Data Fig. 2. Deletion and complementation of HeLa cell lacking 14-3-3 isoform YWHAZ. a,** Expression levels of 14-3-3 isoform YWHAZ (14-3-3ζ) in generated HeLa cell 14-3-3ζ^-/-^ and complemented cell lines. Normalized to WT cells. N = 3-6 independent biological replicates. Mean ± SEM. **, p<0.01; ***, p<0.001. **b,** Protein levels of various 14-3-3 isoforms in WT HeLa and HeLa 14-3-3ζ^-/-^ cell line. Only isoform that is absent in 14-3-3ζ^-/-^ cells is YWHAZ. Western blots using isoform-specific antibodies. **c,** Protein levels of 14-3-3 isoforms in soluble (cytosolic) fraction of WT HeLa or HeLa 14-3-3ζ^-/-^ cells complemented with nothing (-), empty vector, or HA-tagged 14-3-3ζ. Western blot using antibodies that recognize both β and ζ isoforms (top blot) or all 14-3-3 isoforms (middle blot). **d,** Protein levels of 14-3-3ζ in 14-3-3ζ^-/-^ cell line complemented with HA-tagged 14-3-3ζ derivatives defective in 14-3-3ζ dimerization (monomer), in binding phosphorylated ligands (binding pocket), or in interaction with vimentin (vimentin). Western blot using antibodies to HA. β-actin, loading control.

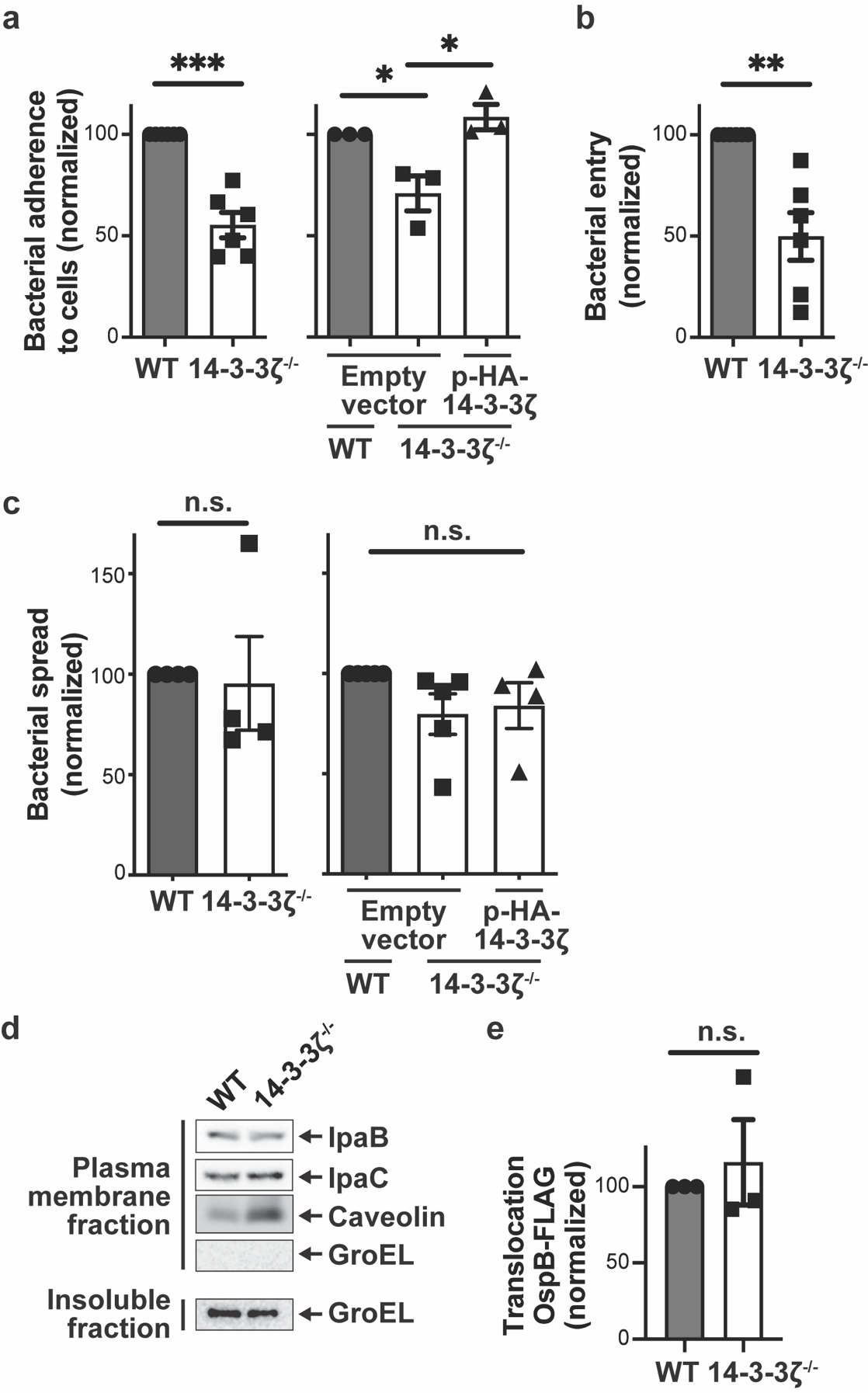

**Supplementary Data Fig. 3.** **14-3-3ζ is required for efficient bacterial adherence to and entry into cells.** **a,** Reduction of *S. flexneri* adherence to HeLa cells lacking 14-3-3ζ, which is rescued by complementation via transfection with HA-tagged 14-3-3ζ. Entry adherence defined as the number of bacteria associated with the monolayer at 30 mins of infection without gentamicin (see Methods). Normalized to WT HeLa (left graph) or WT HeLa complemented with empty vector (right graph). N = 6 (left graph) or 3 (right graph) independent biological replicates. **b,** Reduction of *S. flexneri* entry into HeLa cells lacking 14-3-3ζ, as determined by number of bacteria recovered from the monolayer at 1 hr 30 min of infection (see Methods). Normalized to WT HeLa cells. N = 6 independent biological replicates. **c,** Bacterial spread through the cell monolayer is unaffected by the absence of 14-3-3ζ. Size of bacterial plaques in monolayers of indicated cell lines. Normalized to WT HeLa (left graph) or WT HeLa complemented with empty vector (right graph). N = 3-5 independent biological replicates. **d,** Insertion of *S. flexneri* type 3 translocases IpaB and IpaC into the plasma membrane is unaffected by the absence of 14-3-3ζ. Western blot of IpaB and IpaC in the plasma membrane fraction. Caveolin, membrane protein control. GroEL, bacterial cytoplasmic protein control. **e,** *S. flexneri* type 3 translocation of effectors into the cell is unaffected by the absence of 14-3-3ζ. Quantification by band densitometry of western blots of FLAG-tagged effector OspB in the cytosolic fraction. Normalized to WT cells. N = 3 independent biological replicates. Mean +/- SEM. n.s., not significant. *, p<0.05; **, p<0.01; ***, p<0.001.

**Supplementary Data Table 1. Significant^a^ human hits from Virotrap**

| **WT vs Uninfected** | | | |
| --- | --- | --- | --- |
| **Gene Symbol** | **Uniprot ID** | **p value** | **Log_2_FC** |
| CAP1 | Q01518 | 0.020 | 3.3 |
| CTSV | O60911 | 0.025 | 2.7 |
| HNRNPC | P07910N | 0.026 | -2.4 |
| CAP2 | P40123 | 0.027 | 3.4 |
| KIF5C | O60282 | 0.027 | -2.7 |
| CLTC | Q00610 | 0.033 | 2.2 |
| MVP | Q14764 | 0.033 | 2.7 |
| IMPDH2 | P12268 | 0.033 | -2.6 |
| CAPZB | P47756 | 0.035 | -2.6 |
| LAMP1 | P11279 | 0.043 | 3.3 |
| **Stalled translocon vs Uninfected** | | | |
| **Gene Symbol** | **Uniprot ID** | **p value** | **Log_2_ FC** |
| ACTN1 | P12814 | 0.001 | 5.1 |
| SERPINB1 | P30740 | 0.002 | 4.5 |
| CLU | P10909 | 0.004 | -3.7 |
| YWHAB | P31946 | 0.005 | 3.6 |
| KIF11 | P52732 | 0.005 | -4 |
| ANXA3 | P12429 | 0.008 | 3.6 |
| CCN2 | P29279 | 0.011 | -5 |
| MLH3 | Q9UHC1 | 0.013 | -3.1 |
| SNRPB2 | P08579 | 0.014 | -3.6 |
| PSMD12 | O00232 | 0.015 | 2.6 |
| CSE1L | P55060 | 0.018 | 3.3 |
| RAN | P62826 | 0.021 | 3.2 |
| S100A14 | Q9HCY8 | 0.022 | 2.4 |
| PAFAH1B2 | P68402 | 0.024 | 2.6 |
| SERPINE2 | P07093 | 0.026 | -3.1 |
| CKM | P06732 | 0.033 | -2.6 |
| QPCT | Q16769 | 0.036 | -2.4 |
| CAP2 | P40123 | 0.039 | 3.1 |
| CFI | P05156 | 0.04 | -3.9 |
| C7 | P10643 | 0.04 | -2.5 |
| H2BC12 | O60814 | 0.05 | 2.3 |
| ALOX12B | O75342 | 0.05 | 2.6 |
| SMOC1 | Q9H4F8 | 0.05 | -2.4 |
| KRT78 | H0YI54 | 0.05 | -2.1 |
| IGKV3-20 | P01619 | 0.05 | 2.4 |

^a^p ≤ 0.05

**Extended Data Table 2. Significant^a^ human hits from Virotrap, dependent on 14-3-3ζ.**

^a^p ≤ 0.05

**Extended Data Table 3. Strains and plasmids used in this study.**

| **Strains** | **Genotype or description** | **Reference or source** |
| --- | --- | --- |
| *S. flexneri* 2457T | Wildtype serotype 2a | ^53^ |
| *S. flexneri* BS103 | 2457T cured of its virulence plasmid | ^64^ |
| *S. flexneri* ∆*ospB* | 2457T ∆*ospB* | Gift of Cammie Lesser |
| *S. flexneri* ∆*ipaB* | 2457T ∆*ipaB* | ^7^ |
| S. flexneri ∆*ipaB ipaC* | 2457T ∆*ipaB* ∆*ipaC* | This study |
| *E. coli* sfT3SS | pmT3SSΔeff, pNG162 v*irB,* pBAD33 *pilT*, Spec^r^, Kan^r^, Cm^r^ | Gift of Cammie Lesser ^63^ |
| HeLa CCL2 |  | ATCC |
| HeLa CCL2 YWHAZ^-/-^ |  | This study |
| HeLa CCL2 CAP2^-/-^ |  | This study |
| HeLa CCL2 YWHAZ^-/-^ CAP2^-/-^ |  | This study |
| **Bacterial plasmids** |  |  |
| pIpaB | pDSW206 *ipaB*, Amp^r^ | ^7^ |
| pIpaB-FLAG | pDSW206 i*paB*-FLAG, Amp^r^ | This study |
| pIpaB-FLAG IpaC R362W | pDSW206 *ipaB*-FLAG i*paC* R362W, Amp^r^ | This study |
| pOspB-FLAG | pDSW206 *ospB*-FLAG, Amp^r^ | Gift of Cammie Lesser |
| pAfa1 | pNG162 *afa1*, Spec^r^ | ^56^ |
| pIpaB-FLAG IpaC | pDSW206 i*paB*-FLAG i*paC*, Amp^r^ | This study |
| pmCherry | pBR322 mCherry | Lab stock |
| **Mammalian plasmids** |  |  |
| GAG | pMET7 GAG-GW, | ^18^ |
| FLAG-VSVG | pcDNA3 FLAG-VSVG | ^18^ |
| VSVG | pcDNA3 VSVG | This study |
| Empty | pIRESpuro3 | Addgene |
| HA-WT | pIRESpuro3 HA-YWHAZ | This study |
| Monomer | pIRESpuro3 HA-YWHAZ L_12_AE-->Q_12_QR | This study |
| Binding pocket | pIRESpuro3 HA-YWHAZ K_49_R_56_R_127_Y_128_-->A_49_A_56_A_127_A_128_ | This study |
| Vimentin | pIRESpuro3 HA-YWHAZ Y_82_REKIE-->Q_82_RENIQ | This study |
